# EEG biomarkers of reinforcement learning and motivation: A multi-task battery

**DOI:** 10.64898/2026.06.28.735051

**Authors:** Zeguo Qiu, Muzhi Wang, Haoyang Lu, Yaniv Abir, Raziya Zharmakhan, Nikita Singh, Michaela Poppe, Luigi A.E. Degni, Quentin J. M. Huys

## Abstract

Serotonin and dopamine make dissociable contributions to reinforcement learning (RL) sub-components, yet we lack neural biomarkers capable of detecting their differential effects. Here, we report the development of a five-task EEG battery designed to probe these dissociable RL mechanisms. Using a model-free analytical approach in healthy volunteers, we identify distinct neural markers across probabilistic instrumental learning, motivational vigour, Pavlovian-instrumental transfer, reversal learning, and a working memory-RL task. A centroparietal P300 tracked incremental learning across three paradigms and showed cross-task convergent validity. Readiness potentials and beta suppression indexed value-based motor preparation, while frontal theta captured Pavlovian-instrumental conflict. The largely independent pattern across markers supports the battery’s capacity to detect selective pharmacological effects on distinct neural systems.

## Introduction

Adaptive behaviour depends on a suite of dissociable learning and motivational processes including learning action values from feedback and adjusting the vigour of actions in proportion to reward rates. Reinforcement learning (RL) provides a powerful computational framework for formalising these processes and linking them to their underlying neural and neurochemical substrates (Sutton & Barto, 1998; Dolan & Dayan, 2013; Gershman et al., 2024).

Neuromodulators such as serotonin and dopamine are richly implicated in the neural implementation of RL in the brain. A recent systematic review and meta-analysis of pharmacological studies showed that dopaminergic and serotonergic manipulations have distinct associations with specific RL sub-components: while dopamine was selectively associated with reward learning, reward sensitivity, and action vigour, serotonin was associated with punishment learning and aversive inhibitory processes (Mkrtchian et al., 2025). These findings align with a substantial body of research on the underlying mechanistic processes. For example, phasic dopaminergic signalling has been shown to encode reward prediction errors (Schultz et al., 1997), and tonic dopamine has been linked to the average reward rate that governs action vigour (Niv et al., 2007). Serotonin, by contrast, has been most prominently associated with aversive processing, punishment-based learning, and inhibitory responses to threatening stimuli (Cools et al., 2008; Dayan & Huys, 2009; Malamud et al., 2024), though its role in appetitive contexts is increasingly recognised (Michely et al., 2022; Colwell et al., 2024).

Electroencephalography (EEG) offers a powerful tool to investigate the neural dynamics underlying these neuromodulatory influences on RL processes with millisecond temporal resolution. A rich body of work has characterised EEG signatures across the RL domains described above. In instrumental learning, frontocentral deflections occurring approximately 250-350 ms after outcome feedback (the P3a; feedback-related negativity/FRN; reward positivity/RewP) have been established as neural correlates of reward prediction error signalling (Holroyd & Coles, 2002; Proudfit, 2015). A later centroparietal P300 or P3b component following feedback has been linked to context updating and the integration of outcome information into ongoing value representations (Donchin & Coles, 1988; Twomey et al., 2015).

More recently, pharmacological studies have linked these signatures to specific neuromodulatory systems: amphetamines modulate the RewP during instrumental learning (Cavanagh et al., 2022), noradrenergic agents alter P300 amplitude (Jepma et al., 2018), and serotonergic manipulations affect feedback processing in ways that may be asymmetric across reward and punishment (Michely et al., 2022; Colwell et al., 2024).

In the time-frequency domain, frontal theta oscillations (3–7 Hz) have been consistently implicated in cognitive control and conflict monitoring across a range of paradigms (Cavanagh & Frank, 2014; Cohen, 2014), including Pavlovian-instrumental conflict tasks where the need to override prepotent approach or avoidance tendencies engages medial frontal control processes (Cavanagh et al., 2013). In addition, premovement sensorimotor beta suppression (13–30 Hz) is a well-established marker of motor preparation (Pfurtscheller & da Silva, 1999) that has more recently been shown to reflect motivational modulation of action, with greater suppression preceding actions directed toward higher-value outcomes (Kilavik et al., 2013).

However, most of the existing work has examined individual markers in isolation within single paradigms. The broader question of how EEG signatures across multiple RL domains relate to one another, and whether they provide dissociable targets for pharmacological intervention, remains largely unaddressed.

These questions carry direct clinical relevance. Meta-analyses have documented impairments across instrumental learning, motivational vigour, cognitive flexibility, and working memory contributions to learning in depression (Halahakoon et al., 2020; Pike & Robinson, 2022; Husain & Roiser, 2018; Treadway & Zald, 2011). Given that serotonergic and dopaminergic systems make dissociable contributions to these RL processes (Mkrtchian et al., 2025), it raises the possibility that depressed individuals with different profiles of RL dysfunction may respond differentially to serotonergic versus dopaminergic treatments. However, clinicians currently lack principled methods for selecting among these agents for individual patients, a process that relies largely on trial and error, burdensome for patients and costly for healthcare systems (Maia & Frank, 2017).

Here, we report the development of a comprehensive EEG task battery, as part of the Reinforcement Learning Mechanisms of Antidepressant Treatments (RELMED) project. The motivation of including multiple tasks in our battery is straightforward: single-paradigm studies cannot determine whether an observed pharmacological effect reflects a specific RL mechanism or a more general influence on task performance. A multi-task battery can address this by establishing which neural signatures covary across paradigms, likely indicating a shared underlying process, and which remain independent, allowing effects to be attributed to distinct RL sub-components.

In the present study, we aimed to provide dissociable neural biomarkers across multiple RL subcomponents. Five EEG paradigms were administered to healthy volunteers: probabilistic instrumental learning, a vigour task, Pavlovian-instrumental transfer, reversal learning, and working memory reinforcement learning. For each task, we selected EEG markers based on established links to the targeted RL processes and evaluated their sensitivity to the experimental manipulations most relevant to future pharmacological investigation. Finally, we examined cross-task correlations among these markers to assess convergent and divergent validity across the battery, testing whether the identified signatures index shared or dissociable neural processes.

## General method

### Participants

Healthy adult volunteers participated in the study. All participants had normal or corrected-to-normal vision and no history of neurological or psychiatric disorders. The study was approved by the UCL Research Ethics Committee (project ID: 1273). An overview of each task including sample sizes and trial counts is provided in Table 1.

**Table 1:** Overview of the five EEG tasks, sample sizes, trial counts, and identified EEG markers.

| Task | RL construct measured | N (final EEG) | Total trials | EEG markers identified |
| --- | --- | --- | --- | --- |
| PILT – blocked (Exp. 1a) | Incremental value learning | 30<sup>a</sup> | Up to 240 | P300 (trial progression), Pe (outcome valence), FRN (loss magnitude), P100 (loss magnitude), neural value representation shift (RSA) |
| PILT – mixed valence (Exp. 1b) | Incremental value learning | 32–33<sup>b</sup> | Up to 200 | P300 (trial progression), Pe (outcome valence), P100 (reward and loss magnitude), RSA |
| Vigour (Exp. 2) | Motivational modulation of action vigour by reward rate | 65 | 54 | Readiness potential (reward-per-press) |
| PIT (Exp. 3) | Pavlovian modulation of instrumental behaviour | 63<sup>c</sup> | 108–144 | Beta suppression (Pavlovian valence, instrumental value), frontal theta (aversive conflict) |
| Reversal learning (Exp. 4) | Behavioural adaptation to changing reward contingencies | 51<sup>d</sup> | Variable | Centro-parietal negativity (confirmatory outcomes), FRN (expectation violation), P300 (expectation violation), frontal positivity (stable vs. volatile surprise) |
| WM-RL (Exp. 5) | Dissociation of WM-supported and incremental RL | 44–47<sup>e</sup> | 108 | P300 (stimulus repetition), FRN (reward magnitude), late frontal positivity (reward magnitude) |

<sup>a</sup> 32 recruited; 1 lost due to technical failure, 1 excluded due to low usable epochs.
<sup>b</sup> 34 recruited; 1 excluded for below-chance performance; N differs between outcome-locked (32) and stimulus-locked (33) analyses.
<sup>c</sup> 65 recruited (35 pilot 1, 30 pilot 2); 2 excluded due to low usable epochs.
<sup>d</sup> 54 recruited; 1 lost due to technical failure, 2 excluded due to low usable epochs.
<sup>e</sup> 51 recruited; 4 excluded for below-chance performance, 3 excluded due to low usable epochs; N differs between outcome-locked (44) and stimulus-locked (47) analyses.

### Sequence generation

A key aim of the study design is to develop measurement tools enabling comparison of RL components across individuals. In mechanistic group studies, sequences of stimuli and rewards are typically randomised between individuals to minimise bias at the cost of increased variance. Here, the focus was instead on developing tools which enable comparisons between individuals, and over multiple time-points. This requires a minimisation of variance, while bias is less critical. To minimise variance at the cost of potential bias, sequences were hence pseudorandomised and fixed across individuals.

Prior to finalising the task battery, we conducted Patient and Public Involvement and Engagement (PPIE) sessions with individuals with lived experience of depression. Feedback was gathered through in-person and online focus groups, and covered aspects including task instructions, response button usability, trial sequence, and overall session length. The feedback directly informed revisions to the battery prior to data collection.

### Apparatus

All stimuli were presented on a monitor with the resolution of 1920 ×1080 pixels. All experimental tasks were coded and run in PsychoPy3 (version 2022.2.5; Peirce et al., 2019). Stimuli were presented on a dark grey background on the monitor screen.

### EEG data acquisition and preprocessing

Continuous EEG was recorded at 1000 Hz using the BrainProducts BrainAmp system or the BrainProducts Actichamp Plus system with 64 active electrodes positioned according to the international 10-20 system. The data were online-referenced to the FCz electrode. Horizontal electrooculogram (EOG) was recorded using a pair of bipolar electrodes placed at the outer canthi.

Pre-processing was conducted using a combination of EEGLAB 2024.1 (Delorme & Makeig, 2004) and ERPLAB (Lopez-Calderon & Luck, 2014) in MATLAB. Bad EEG channels were visually inspected and interpolated using spherical spline interpolation, with the specific channels interpolated varying by participant. EEG data were then down-sampled to 500 Hz, band-pass filtered offline (0.1 - 30Hz for ERP/GLM analyses and 0.1 - 50Hz for Event-Related Spectral Perturbation or ERSP analyses), and re-referenced to the average of all scalp electrodes, excluding the EOG channel. Data segmentation was subsequently performed, with signals time-locked to events of interest in each task.

Independent Component Analysis (ICA) was then performed using the extended infomax algorithm. Components corresponding to eye movements and muscle artifacts were automatically identified using ICLabel (Pion-Tonachini et al., 2019) and rejected when the combined probability of eye- and muscle-related activity exceeded 0.85.

Residual artifacts were further identified using two semi-automated rejection procedures. First, any epochs exceeding a voltage threshold of ±80 µV at any channel were flagged and removed. Second, a moving window peak-to-peak threshold procedure (200 ms window, 100 ms step) was applied to detect high-frequency noise, again using a ±80 µV threshold.

## Experiment 1a: Probabilistic instrumental learning task (blocked valence design)

The probabilistic instrumental learning task (PILT) measures incremental value learning from trial-by-trial feedback (Pessiglione et al., 2006; Halahakoon et al., 2024). Within the RL framework, outcomes drive value estimates via prediction errors, with reward and punishment learning thought to depend on dissociable neuromodulatory systems: dopamine has been linked to reward prediction errors and reward learning, while serotonin is more prominently associated with punishment processing (Dayan & Huys, 2009; Mkrtchian et al., 2025). PILT tasks therefore provide probably the most standard measure of value-based learning with separable windows for probing dopaminergic and serotonergic contributions to RL. The current PILT task incorporates several details worth noting. First, it involves two-alternative forced choices between different visual stimuli, where the two stimuli on each choice are always drawn from two different categories. The task is structured in terms of a series of blocks, with new stimuli for each block. Second, to ensure acceptability and ensure valid measurements from as many participants as possible, the task is built on a shaping protocol, with choices becoming more difficult over time. In the early blocks, outcomes are deterministic, enabling fast learning. In the later blocks, outcomes are increasingly probabilistic. Third, to minimise the duration, blocks can end early if participants choose the better stimulus consistently. Fourth, the task aimed to disentangle outcome sensitivity from learning. To achieve this, multiple different outcome magnitudes were used, across both positive and negative valence.

### Method

#### Stimuli

Object images were obtained and adapted from the “Massive Memory” object categories database (Konkle et al., 2010). Images from a total of 24 categories were used. Each image was 300-by-300 pixels in shape. The luminance of all the object images was matched by using the SHINE-color toolbox in MATLAB (Dal Ben, 2023). Coin images were sourced online and edited using Adobe Photoshop (version 26.8.1, 2025; Adobe Systems, San Jose, CA) to render broken coins.

As will be detailed in the Procedure below, before the presentation of the outcome in each trial, participants saw an outcome-paired Pavlovian stimulus. These Pavlovian stimuli were generated using R 4.4.1 and the ggplot2 package (version 3.5.1). Importantly, for each experimental session, we selected distinctive pairs of contrasting colours and symbols to minimise potential generalisation across sessions. Pavlovian stimuli were divided into appetitive and aversive groups, reflecting opposing valences. To maximise differentiation, we used clearly distinguishable symbols and hues separated by ∼180^◦^ in HCL colour space, while holding luminance (70) and chroma (85) constant. Within each valence, stimuli shared the same colour and symbols, but varied in size and spatial density according to Pavlovian value. Larger absolute coin values (e.g., £1 coin and broken £1 coin) were represented by larger, more sparsely distributed symbols, whereas smaller absolute values (e.g., 1p coin and broken 1p coin) were depicted with smaller, more densely arranged symbols (Figure1A).

#### Procedure

Participants were instructed to learn which stimulus in the pair led to more favourable outcomes through trial and error, and to maximise their accumulated gains. The task consisted of 24 blocks (12 reward blocks and 12 punishment blocks). In reward blocks, participants received either an optimal outcome (a fifty-pence coin or a one-pound coin), or a suboptimal outcome (i.e., one-penny coin). In punishment blocks, the outcomes reflected monetary losses (breaking either a one-penny coin, a fifty-pence coin or a one-pound coin). Within each block, there were a maximum of 10 trials. Importantly, a block could terminate early if participants consecutively chose the optimal stimulus five times.

An important consideration is to ensure that all participants are able to perform the task. To facilitate this, we employed a shaping procedure and varied the feedback structure across blocks. Specifically, the first four blocks were deterministic, such that the optimal stimulus always led to an optimal outcome (100% probability). For block 5 and 6, the optimal stimulus yielded an optimal outcome with a probability of 90%, and a probability of 80% for all subsequent blocks, while the suboptimal stimulus had a low probability (10% or 20%, respectively) of delivering a favourable outcome.

As shown in Figure 1A, each trial started with an intertrial fixation screen for a random duration between 500 and 800ms, after which a pair of stimuli appeared. Participants had up to three seconds to make a choice by pressing the left or right arrow key. Upon a choice of the stimulus, a Pavlovian stimulus, which is deterministically associated with the subsequent outcome, replaced the chosen stimulus and was presented for one second. Afterwards, the outcome associated with the selection was revealed for one second, which ended the trial. A practice block consisting of four trials were provided prior to the main task.

**Figure 1:**
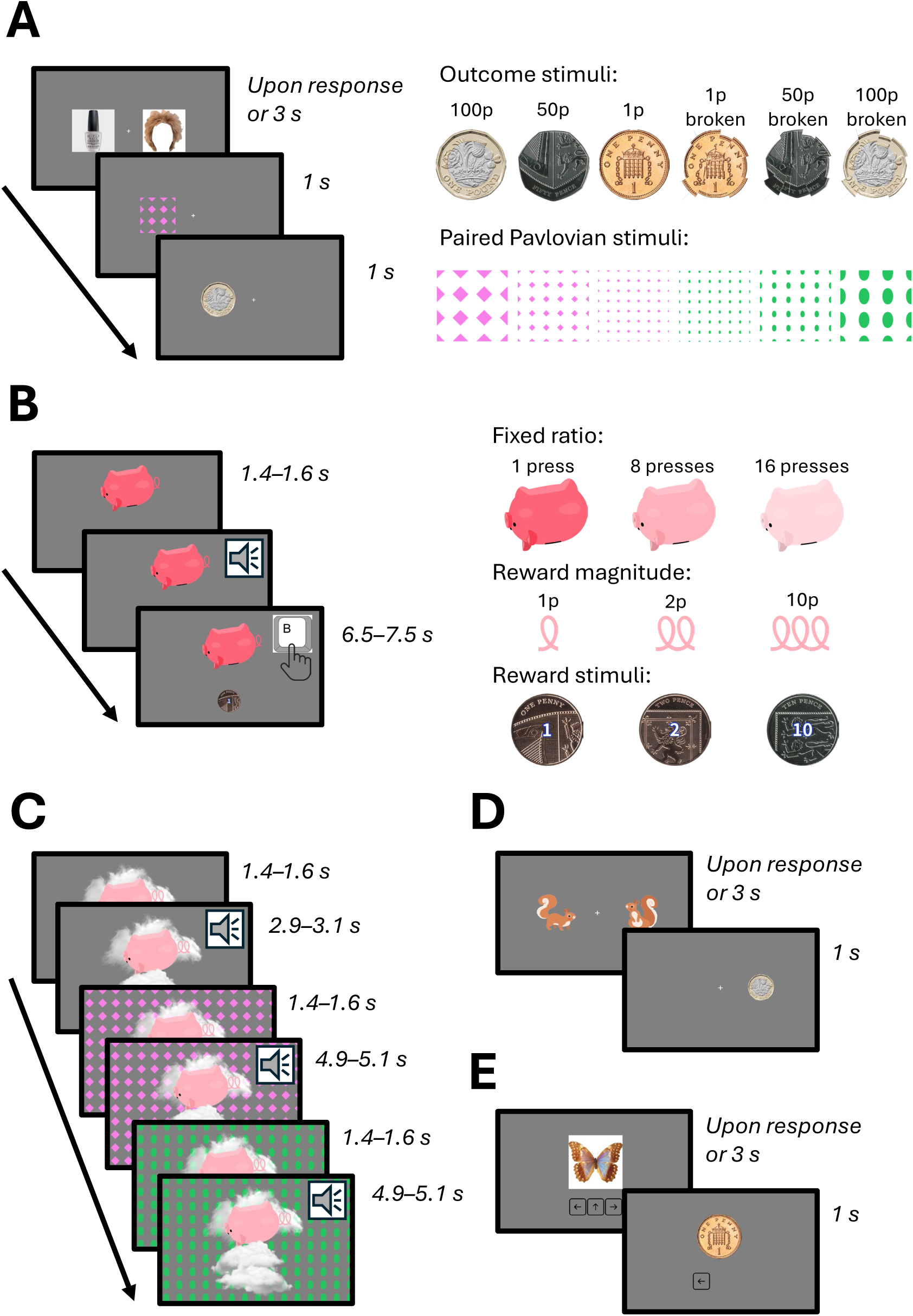
(A) Time-course of events and stimuli during a trial of Experiment 1a and 1b (PILT), in which participants learned to choose between two stimuli to maximise monetary outcomes via trial-and-error feedback. (B) Time-course of events and stimuli during a trial of Experiment 2 (Vigour task), in which participants pressed a key repeatedly to earn rewards, with effort requirements and reward magnitudes varying across trials. (C) Time-course of events during a trial of Experiment 3 (PIT task), in which Pavlovian background cues of varying affective valence were presented during instrumental responding. (D) Time-course of events during a trial of Experiment 4 (Reversal learning task), in which participants tracked which of two stimuli was currently optimal as reward contingencies periodically reversed. (E) Time-course of events during a trial of Experiment 5 (Working memory task), in which participants learned stimulus-action mappings under varying delays between stimulus repetitions.

Additionally, in order to probe a potential neural representation shift induced by probabilistic instrumental learning, participants completed an incidental viewing task before and after the probabilistic instrumental learning task. In this task, the stimuli used in the main learning task were presented centrally on the screen, one after another. Each stimulus was presented for 1s and repeated four times throughout each incidental viewing session. An intertrial interval (0.8–1.4s, uniformly distributed) was presented between stimulus presentation with a fixation cross remained visible throughout. To ensure sustained attention, on 40 out of the total of 192 trials, the fixation cross changed colour for 500ms, and participants were required to press the spacebar when they detected this change.

#### EEG pre-processing and analysis

We epoched the data time-locked to the onset of object images or the onset of outcome images, depending on the analysis, using the interval between −200 and 0 ms before the onset of events (object images or outcomes) as the baseline for correction.

We used a mass-univariate general linear model (GLM) to analyse trial-wise EEG. For each participant and time-locked event, an ordinary least squares (OLS) GLM with an intercept was fit at every electrode and time sample using trial-level behavioural regressors z-scored within participant. Specifically, a stimulus-locked GLM was run to test whether EEG amplitudes encoded learning, as indexed by trial progression (controlling for side of the stimulus). An outcome-locked valence-by-magnitude GLM was run to test main effects of outcome valence and outcome magnitude and their interaction. For each participant, *β*-coefficients were obtained at all electrodes and time points, yielding 3-D *β*-maps. Group-level inference tested *β* against zero using two-sided sign-flip permutations across subjects (5,000 permutations) with Threshold-Free Cluster Enhancement (TFCE) over all channels and timepoints within an epoch, controlling family-wise error at *α* = .05 via the max-TFCE statistic. Analyses were implemented in customised MATLAB code with FieldTrip v2024a.

#### Representation similarity analysis

To examine how value representations evolve over the course of learning, we conducted a time-resolved representational similarity analysis (RSA) comparing neural activity patterns to model-based value predictions.

First, EEG data were downsampled to 250 Hz to reduce computational demands. Neural representational dissimilarity matrices (RDMs) were constructed by averaging epoched data within each stimulus type separately for pre- and post-learning sessions. For each time point, we extracted neural activity patterns spanning a 60-ms temporal window (15 consecutive time points × 64 channels), which served as multidimensional feature vectors for each stimulus. Pairwise Euclidean distances between these feature vectors were computed to construct a neural RDM at each time point, yielding a time-resolved series of neural RDMs across the entire epoch.

Model RDMs were derived from the learned stimulus values during the learning phase. For each participant and each stimulus, we computed the mean outcome value across all trials in which that stimulus was chosen. A model RDM was then constructed by computing pairwise absolute differences in these mean values across all stimulus pairs, reflecting the dissimilarity structure of learned stimulus values.

At each time point and for each participant, we quantified the correspondence between neural and model RDMs using Kendall’s tau correlation coefficient. To assess learning-related changes in value representations, we computed the difference in correlation strength between post-learning and pre-learning sessions for each time point.

Group-level statistical inference was performed using the same TFCE-based cluster permutation testing approach as described for the ERP analyses (5,000 permutations, two-sided sign-flip test, family-wise error rate *α* = .05). All RSA computations were implemented in Python using the MNE-RSA package.

### Results

#### Behaviour

Behavioural results demonstrated robust reinforcement learning, with participants more likely to select the correct choice as previous correct trials accumulated (logistic regression: *β* = 0.37, z = 14.95, *p < .*001; OR = 1.45, 95% CI: [1.38, 1.52]). Learning curves are separately plotted for reward and punishment blocks in Figure 2A.

**Figure 2:**
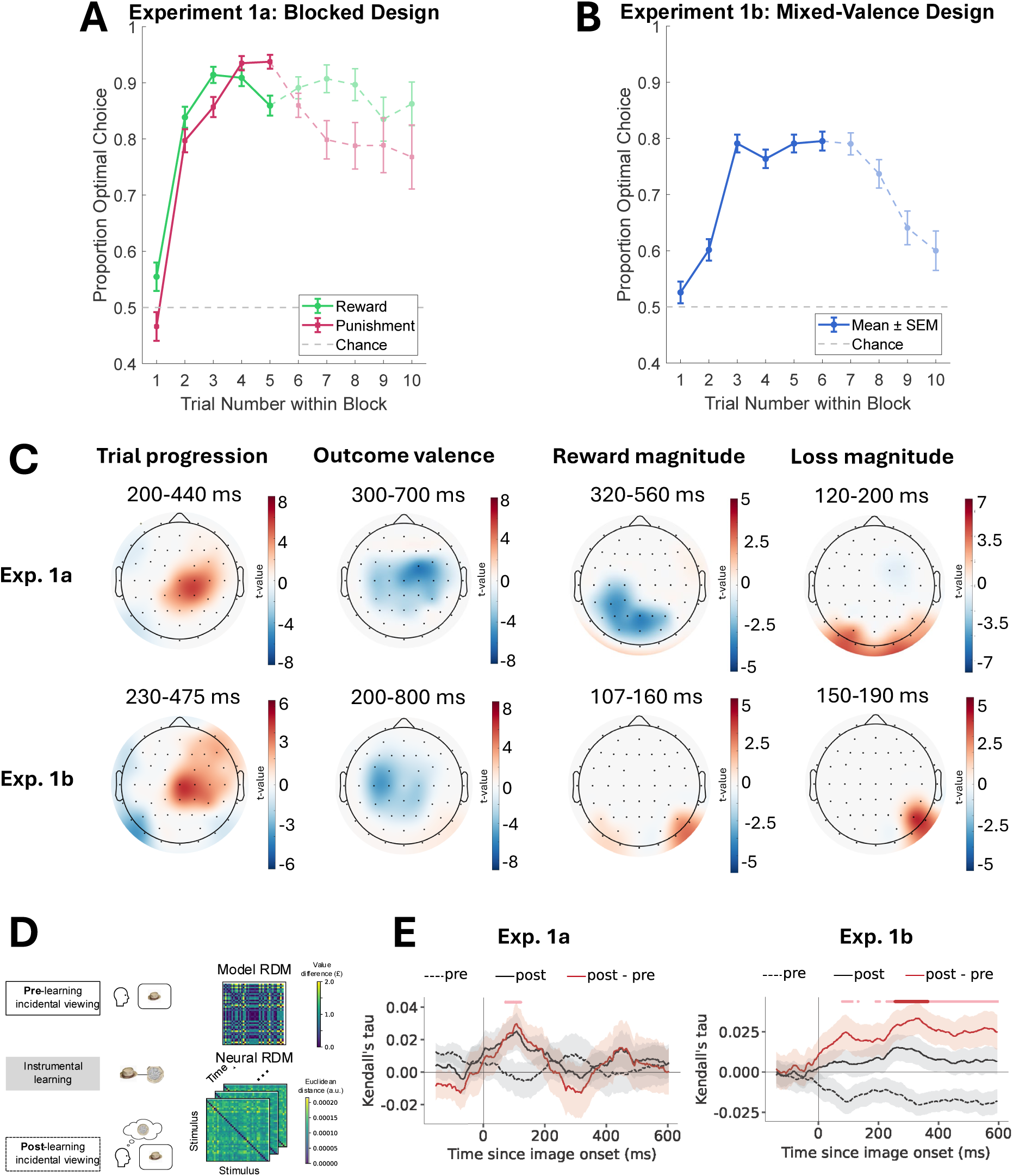
(A) Learning curves for the reward and punishment learning blocks in Experiment 1a (blocked design). (B) Learning curve for Experiment 1b (mixed-valence design). Dashed lines indicate trial positions that were reached by fewer participants due to early block termination following five consecutive optimal choices. (C) Scalp topography plotting significance-thresholded average regression weights for trial number within each block (controlling for side of the stimulus), outcome valence, magnitude of the rewarding feedback and punishing feed-back, for Experiment 1a and 1b. White is nonsignificant; warm colours are positive regression weights; cold colours are negative regression weights. (D) Procedure of the RSA and schematic diagram of the model and neural RDMs. (E) Time course of Kendall’s tau between neural and model RDMs, for pre- and post-learning stimulus representation and their difference in Experiment 1a and 1b. The horizontal aligned dots denotes significance of *t* test between Kendall’s tau between post and pre (pink, uncorrected *p <* 0.05; red, TFCE-corrected *p <* 0.05)

#### Neural signatures of learning

We first examined how stimulus-locked EEG signals varied as a function of the trial progression, using a general linear model (GLM). A significant centroparietal cluster (spatial peak: CP2) from 200-440 ms post-stimulus (temporal peak: 323 ms) was found, whereby later trials were associated with a stronger P300 than earlier trials, *t*(29) = 7.69, TFCE-corrected *p < .*05 (Figure 2A, left).

#### Neural signatures of outcome evaluation

To assess neural activity during outcome evaluation, we examined EEG responses to feedback stimuli using a GLM with separate magnitude slopes for positive and negative valence trials. The model included valence as a main effect (+1/-1 coding) and z-scored magnitude effects separately gated by trial valence (Mag—Pos, Mag—Neg). A sustained frontocentral negativity was observed for rewards compared to losses between 300 and 700 ms (*ts >* 2.02, TFCE-corrected *ps < .*05; Figure 2C, second column), which reflected a pronounced error-related activation over the parietal regions (Pe; Falkenstein et al., 2000).

Outcome magnitude modulated distinct ERP components depending on valence context. In the reward blocks, a centroparietal P300 was enhanced by a smaller reward, relative to a larger reward (spatial peak: CP3, temporal peak: 419 ms), *t*(29) = 4.74, TFCE-corrected *p < .*05). In the punishment blocks, a larger loss (£1 loss) is associated with significantly more positive ERP amplitudes between 120 and 200ms over posterior electrodes than a smaller loss (i.e., 1-penny loss), with a spatial peak at O2 and a temporal peak at 153ms *t*(29) = 6.68, TFCE-corrected *p < .*05. That is, we observed a valence-specific asymmetry: smaller rewards were associated with enhanced P300 amplitudes, whereas larger losses produced stronger P100 responses than smaller losses. The enhanced P100 may suggest greater attentional capture by severe negative outcomes. By contrast, it is likely that frequency contributes to the effects in the reward blocks, where the smaller rewards are also more surprising or infrequent, thereby eliciting a stronger P300 despite their lower value.

In addition, we found a significant feedback-related negativity towards larger than smaller losses over frontal electrodes between 170 and 200ms (spatial peak: FC2, temporal peak: 181ms; *t*(29) = 4.18, TFCE-corrected *p < .*05), consistent with previous literature on the modulatory effect of negative outcomes on frontal neural activity (Holroyd & Coles, 2002). Note that this FRN was only present in the punishment blocks.

#### Learning alters neural representation

To examine whether instrumental learning alters automatic neural representations of task stimuli, we compared representational patterns before and after the learning phase. Specifically, we tested whether exposure to reward and punishment contingencies during the probabilistic instrumental learning task led participants to represent stimuli of similar associated value as more similar upon mere viewing.

We employed RSA to quantify these changes. For each time point, we constructed representational dissimilarity matrices (RDMs) based on multivariate EEG activity patterns across electrodes within short temporal windows. We then assessed the correspondence between neural dissimilarity and the absolute value difference between stimuli, yielding a time-resolved measure of value representation strength (Figure 2D). This approach allowed us to track the temporal dynamics of value encoding in passive viewing tasks conducted before and after learning (Figure 2E).

We did not observe a significant difference in value-related representational similarity between pre- and post-learning sessions after correction. An uncorrected effect emerged in an early time window (64–100 ms; uncorrected *ps < .*05), but this did not survive correction for multiple comparisons (TFCE-corrected peak *p* = .24, peak *t* = 2.37). One possible explanation lies in the blocked structure of Experiment 1a itself. Because reward and punishment outcomes were presented in separate blocks, stimuli of the same valence necessarily co-occurred within a block, while stimuli of differing valence rarely shared the same temporal context. This blocked exposure may have introduced a valence-context structure superimposed on stimulus value, such that neural similarity between stimuli was shaped not only by their learned outcome value but also by block membership. As our model RDM was constructed solely from the absolute difference in mean stimulus value, it may not have adequately captured this additional valence-context structure, potentially obscuring any genuine value-related representational shift. We return to this issue in Experiment 1b, where reward and punishment outcomes were interleaved on a trial-by-trial basis, removing the confound between value learning and block-level valence context.

## Experiment 1b: Probabilistic instrumental learning task (mixed-valence design)

In Experiment 1a, because reward and punishment contexts were blocked, participants could adopt block-specific shortcuts (e.g., response biases) early in a block. To better capture trial-and-error learning under uncertainty, Experiment 1b interleaved reward and punishment outcomes on a trial-by-trial basis, preventing advance knowledge of the current valence context and discouraging strategy carry-over. This version therefore imposes genuine valence uncertainty, allowing us to assess whether the behavioural and neural signatures of value learning remain consistent under more naturalistic learning conditions.

### Method

#### Stimuli

Object images were chosen from the same picture database and were perceptually matched separately for this experiment.

#### Procedure

Similar to Experiment 1a, participants were instructed to maximise their accumulated gains over the experiment. A block may terminate early given five consecutive choices of the optimal stimulus were made by the participants. Within each of the 20 blocks, the valence of the stimuli-associated outcomes was intermixed. That is, a stimulus in a given pair could result in a reward (obtaining a coin), while the other stimulus in the same pair could result in either reward or punishment (losing a coin). A shaping protocol was implemented such that blocks 1-4 were fully deterministic, while blocks 5-8 and blocks 9-20 were probabilistic, with favourable outcomes delivered on 90% and 80% of trials, respectively. The procedural structure of a trial was the same as in Experiment 1a.

In addition to the main learning task, participants also completed incidental viewings of the stimuli shown in the PILT before and after the task. The viewing session consisted of 160 trials in total (4 repetitions of 40 stimuli). To ensure sustained attention of the participants, they were asked to perform a fixation colour change detection task (32 colour change trials).

#### EEG analysis

The EEG analyses were the same as those in Experiment 1a.

### Results

#### Behaviour

Behavioural results demonstrated robust reinforcement learning, with participants more likely to select the correct choice as previous correct trials accumulated (*β* = 0.38, z = 18.24, *p < .*001; OR = 1.47, 95% CI: [1.41, 1.53], see the learning curve in Figure 2B).

#### Neural signatures of learning

The trial progression GLM revealed that later trials were associated with an enhanced P300 between 230 and 475 ms post-stimulus (spatial peak: CP2, temporal peak: 327 ms), *t*(32) = 6.23, TFCE-corrected *p < .*05. This result showed that trial progression or stimulus learning was likely reflected in centroparietal activity, consistent with the P300 component indexing value-based evidence accumulation.

#### Neural signatures of outcome evaluation

An outcome-locked GLM examining the effects of outcome magnitude and valence revealed a similar valence main effect on the EEG signals, whereby we observed a sustained frontocentral negativity for rewards compared to losses between 200 and 800 ms (*ts >* 2.01, TFCE-corrected *ps < .*05; Figure 2C). In addition, between 180-260ms, there was also a significant posterior positivity towards reward outcomes, compared to losses (spatial peak: P8), *t*(31) = 8.86, TFCE-corrected *p < .*05.

Different from Experiment 1a, we did not observe a P300 towards smaller rewards. This is likely due to the more balanced occurrence probabilities for the lower- and higher-valued rewards. Specifically, as the 1-pence coins are less infrequent in this version of task, this particular feedback is not as surprising as in Experiment 1a, removing its potential oddball-like effect on the P300.

Early posterior components were sensitive to outcome magnitude in both valence contexts. For losses, the P100 was enhanced in response to larger relative to smaller losses (spatial peak: P6) between 150 and 190ms, *t*(31) = 5.13, TFCE-corrected *p < .*05. For rewards, the P100 was also enhanced for larger relative to smaller rewards (spatial peak: O1) between 107 and 160ms, *t*(31) = 4.70, TFCE-corrected *p < .*05. Unlike Experiment 1a (the blocked version) where early magnitude effects were specific to punishment, the mixed design revealed symmetric magnitude sensitivity across both valences, suggesting that trial-by-trial valence uncertainty may enhance early attentional processing of outcome magnitude regardless of valence.

#### Neural representation shift after learning

To assess whether the mixed-valence learning context similarly induced a shift in neural representations, we applied the same RSA pipeline as in Experiment 1a.

Consistent with Experiment 1a, we found that learning successfully altered the neural representation of the stimuli. However, the temporal dynamics of this shift differed markedly from the blocked design. While Experiment 1a showed early perceptual modulation, the mixed-valence conditioning in Experiment 1b was associated with a later emergence of value encoding. Specifically, post-learning EEG patterns (as opposed to pre-learning EEG) showed significantly higher similarity to the value difference matrix in a late time window (Figure 2E; 294–344 ms; *ts >* 3.33, TFCE-corrected *ps < .*05, min *p* = .011).

Collectively, the findings from the two experiments support the conclusion that learning shapes the neural representation of task-relevant stimuli, but the temporal locus of this effect is flexible and sensitive to the uncertainty of the learning environment. This result suggests that while instrumental learning consistently modifies the neural representation of stimuli, the nature of this modification depends on the learning context. In the blocked design, value associations may be incorporated into rapid, sensory-motor processing stages. In contrast, the unpredictable valence context of the mixed design appears to delay this integration to later processing stages, potentially reflecting higher-level cognitive evaluation rather than early sensory-motor modulation.

#### Test-retest reliability of the neural markers

Twenty-two participants returned to the lab for a retest session (range of days between session 1 and 2: 11-19) for the mixed-valence PILT. For the neural signatures reported above, we extracted the mean betas of the significant clusters from session 1, separately for positive and negative clusters. Then, we extracted the means of the same clusters in data from session 2, and performed a Pearson’s correlation between session 1 and 2 on the cluster means for each neural marker. While the correlation was low for the reward magnitude effect between sessions (r = 0.26), the other effects were moderately reliable: loss magnitude (P100, r = 0.46), trial progression (P300, r = 0.54), and the positive and negative valence clusters (r = 0.60 and r = 0.61, respectively).

## Experiment 2: Action vigour

The vigour task assesses how action value modulates the speed and vigour of responding. Within RL theory, action vigour reflects the relationship between the average reward rate of the environment, which is thought to be encoded by tonic dopamine, and the cost of acting quickly (Niv et al., 2007). This task used a fixed-ratio setup, independently manipulating reward magnitude (the amount of reward obtained) and effort requirement (the number of button presses per reward), yielding a reward-per-press proxy for subjective action value (Figure 1B). Motivational deficits in depression are hypothesised to reflect disrupted dopaminergic signalling of opportunity cost (Treadway et al., 2012; Husain & Roiser, 2018), making this task a targeted assay of this mechanism. Several design features are worth noting here. Stimuli were presented on the screen for 1.4-1.6s before an auditory tone indicated participants could start pressing. This period is akin to the decision time in tasks asking participants to decide between options where reward are discounted by different amounts of effort (Treadway et al., 2015; Gold et al., 2013; Berwian et al., 2020; Hewitt et al., 2025). While these designs have repeatedly shown relationshps to anhedonia and depression, the interpretation is complex, with effort, time and uncertainty confounded. After the auditory tone, participants had 6.5-7.5s to press and obtain as many rewards as possible. This period enables a more directly assessment of the opportunity cost of time (Nair et al., 2020). Finally, as we will see below, this design facilitated an efficient assessment of Pavlovian-Instrumental Transfer effects.

### Method

#### Stimuli

The piggy bank stimulus was custom-designed and created in Adobe Illustrator 2025. Saturation was manipulated relative to a baseline image (fixed-ratio 8): for fixed-ratio 1, saturation was doubled (multiplied by 2), for fixed-ratio 8 it was unchanged, and for fixed-ratio 16 it was halved (multiplied by 0.5), see Figure 1B. These adjustments were applied by scaling the saturation channel in HSV space in MATLAB, with out-of-range values clipped at 1. The same adjustment was also performed on the piggy tail baseline image, to render piggy tails at fixed-ratio 1 (doubled saturation), 8 (unchanged), and 16 (halved saturation).

The tail length of the piggy bank stimulus varied according to trial reward magnitude, with longer tails indicating higher reward values. Coin images (one penny, two pence, and 10 pence) were sourced online and standardised for size consistency across experimental trials.

#### Procedure

Each trial began with the presentation of a piggy bank stimulus (see Figure 1B), followed by a jittered preparatory interval (1.4–1.6 s). Participants were instructed to withhold responding during this interval. The interval would restart if a premature response was made. A brief auditory cue (500 Hz, 100 ms) then signalled the start of the response phase, which lasted 6.5–7.5 s (jittered).

During the response phase, participants pressed the “B” key with their right index finger to “shake” the piggy bank. Each piggy bank was associated with a specific reward magnitude (1p, 2p, or 10p coin) and effort requirement (fixed ratios of 1, 8, or 16 presses per coin). Successful completion of the ratio requirement triggered a coin animation, with coins dropping from the piggy bank on the screen.

The task comprised one block of 54 trials, covering all 9 reward magnitude × effort combinations in pseudo-random order. Before starting the main task, participants completed a short practice phase in which they familiarised themselves with the piggy bank animation and coin drops.

#### EEG pre-processing and analysis

Epochs were extracted from −1500 to +200 ms relative to the first motor action in each trial, with a pre-epoch baseline of −1500 to −1000 ms.

Given the low number of trials, we pre-defined a region of interest and focused on ERP analyses. Specifically, we examined the readiness potential, a marker of motor preparation (Deecke, 1996; Fried et al., 2011), at electrode Cz. For each trial, the mean amplitude was extracted between −500 and 0 ms relative to the first motor action. The main predictor of interest was reward-per-press (RPP), computed as reward magnitude divided by required presses. Trial-level mean amplitudes and corresponding RPP values were concatenated across participants and analysed using a linear mixed-effects model in MATLAB, with subject included as a random intercept.

### Results

#### Behaviour

We examined whether RPP predicted participants’ key-press rate (presses/s). A linear mixed-effects model with subject as a random intercept and random slope (*N* = 65) for RPP provided a significantly better fit than a random-intercept-only model (Δ*χ*^2^(2) = 89.38, *p < .*001). In this model, we found that increases in RPP were associated with significantly higher key-press rates (*β* = 0.08, SE = 0.01, *t*(3508) = 9.11, 95% CI: [0.06, 0.10], Figure 3A). The behavioural RPP effect showed moderate test-retest reliability (r = 0.58), confirming that the task captures stable incentive-driven motor invigoration.

**Figure 3:**
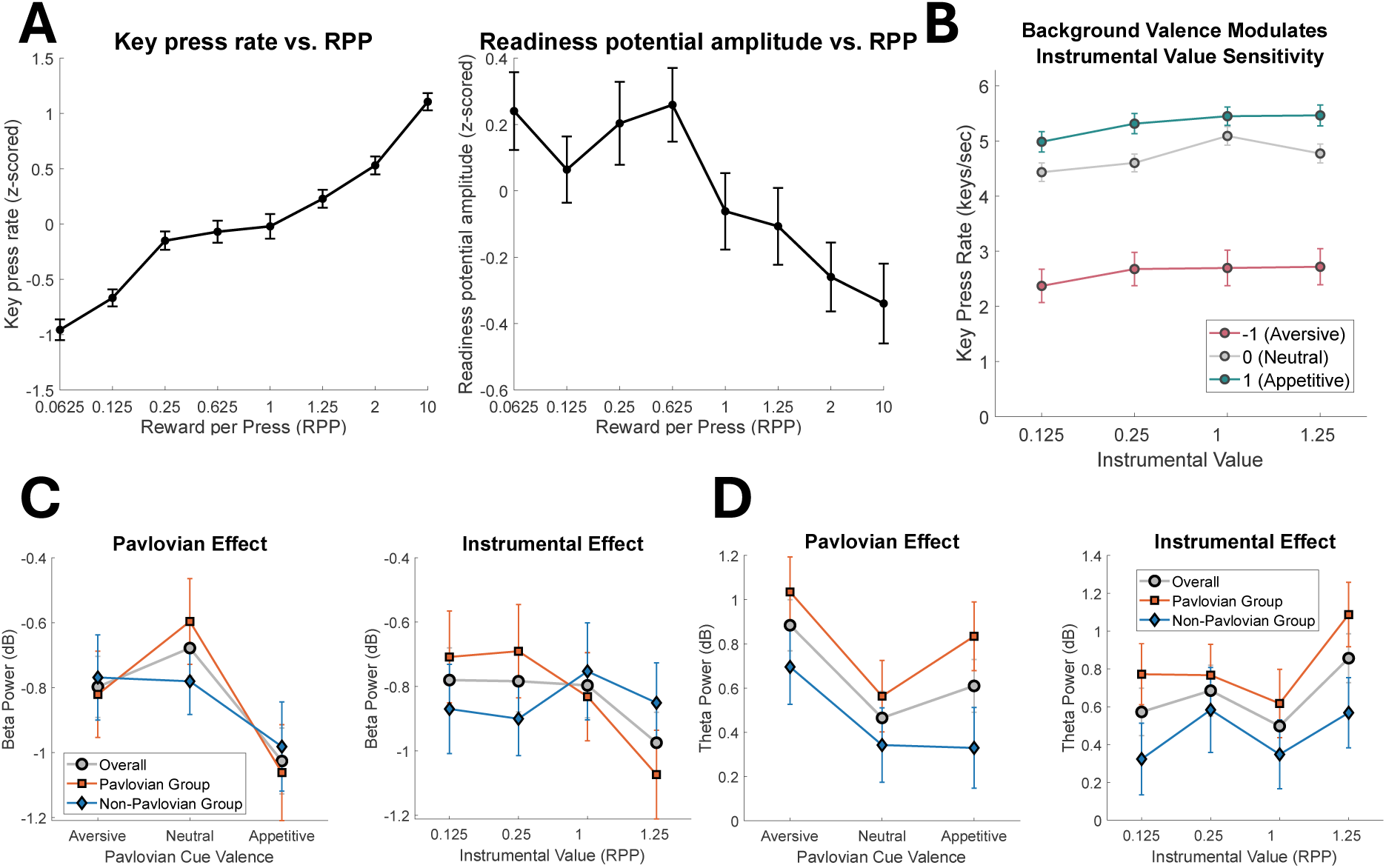
(A) Key press rate increases as stimulus value (reward per press) increases in vigour task. Readiness potential prior to movement decreases as stimulus value increases in vigour task. (B) Key press rate by the background value and instrumental conditions across all participants in the PIT task. (C) Beta power over FC3 by Pavlovian background and instrumental conditions, separated by group. (D) Theta power over Fz by P Pavlovian background and instrumental conditions, separated by group.

#### Readiness potential scales with the subjective value of actions

The linear mixed effect model revealed that RPP significantly predicted readiness potential amplitude (*F*(1, 3451) = 7.86, *p* = .005) whereby higher RPP values were associated with more negative pre-motor activity at Cz (readiness potential: *β* = −0.17 µV, SE = 0.06, 95% CI: [-0.29, -0.05], Figure 3A). This suggests that, in this task, participants showed stronger motor preparation when actions were subjectively more valuable. The readiness potential showed robust group level modulation by incentive value in session 2 as well (readiness potential: *β* = −0.23 µV, SE = 0.10, 95% CI: [-0.42, -0.03]), but individual-difference reliability could not be meaningfully assessed in the current dataset. With only ∼ 6 trials per condition per subject, single-subject readiness potential estimates carry substantial measurement error. This is a well-known constraint in ERP research: reliable individual-difference measurement requires considerably more trials than detecting group-level effects. Though this does not imply that neural modulation is inherently unstable. Rather, the current design was optimised for detecting within-subject experimental effects across conditions, not for quantifying between-subject trait-like variation.

## Experiment 3: Pavlovian-instrumental transfer

Affective value learning often distinguishes between Pavlovian learning about stimulus values, and instrumental learning about stimulus-action values (Dayan et al., 2006; Huys et al., 2014). While the PILT probed instrumental learning, and the vigour task probed instrumental components of motivation, neither task enables a direct assessment of Pavlovian learning. Pavlovian influences are of substantial relevance given their rich shaping of instrumental behaviour. Reward-predicting cues, for instance invigorate approach behaviour while punishment-predicting cues suppress it, even when causally irrelevant to the current instrumental contingency (Lovibond, 1983; Corbit et al., 2007; Dayan et al., 2006; Huys et al., 2011; Guitart-Masip et al., 2012). When aversive Pavlovian cues accompany rewarding instrumental contingencies, the two systems conflict, requiring active cognitive control to resolve this conflict, a process in which serotonergic systems are particularly implicated (Dayan & Huys, 2009; Colwell et al., 2024).

Building on the preceding vigour task, in our Pavlovian-instrumental transfer task (PIT), participants continued performing the same key-pressing task, but now the monetary outcomes (the coins dropping) were hidden, and Pavlovian (appetitive or aversive) backgrounds or no background (neutral) were displayed during responding (Figure 1C). The Pavlovian stimuli used in this task were conditioned in the preceding PILT, ensuring their affective values were well-established prior to the transfer phase. Of note, after an initial period showing the instrumental (piggy bank) cue only, there were two periods with Pavlovian cues, enabling a direct contrast between instrumental and Pavlovian cues, and between different Pavlovian cues. Work in Pavlovian-instrumental transfer distinguishes between outcome-specific and outcome-general PIT (Balleine, 1994). Building on previous work in clinical samples, we here opted for a version that does not distinguish these components (Garbusow et al., 2016; Chen et al., 2023; Schad et al., 2020).

### Method

#### Stimuli

The piggy bank and tail stimuli were the same as those used in the vigour task. In addition, cloud images were sourced online to occlude instrumental outcomes, with Pavlovian stimulus patterns from Experiment 1 (PILT) tiled across the background. Participants were reminded of the associative values of these Pavlovian stimuli during instructions.

#### Procedure

Each trial began with the presentation of a piggy bank stimulus, followed by a jittered preparatory interval (1.4–1.6 s). Participants were instructed to withhold responding during this interval. A brief auditory tone (500 Hz, 100 ms) signalled the start of the response phase, which lasted 2.9–3.1 s (jittered) on trials without Pavlovian cues and 4.9–5.1 s (jittered) when a Pavlovian cue was present, see Figure 1C. Different from the vigour task, during the response phase, coin animations were hidden behind clouds, but successful completion of the ratio requirement still resulted in coins being collected.

To construct the PIT sequence, four piggy banks from the vigour task were selected (corresponding to 1p at fixed-ratio-1, 2p at fixed-ratio-8, 2p at fixed-ratio-16, and 10p at fixed-ratio-8). Each piggy bank was presented three times in a row: once without a background and twice with a Pavlovian background, making in total 144 (pilot 1) or 108 trials (pilot 2). On Pavlovian trials, a background image was displayed simultaneously with the piggy bank. These backgrounds represented appetitive or aversive cues, which are associated with either gains (+1p, +50p, or +£1) or losses (−1p, −50p, or −£1). Importantly, the reward/loss contingency associated with the Pavlovian cue applied only once per trial and did not scale with the number of key presses. The instrumental contingency of the piggy bank remained unchanged.

#### EEG pre-processing and analysis

Similar to the vigour task, we examined the readiness potential at electrode Cz. For each trial, the mean amplitude was extracted between −500 and 0 ms relative to the first motor action. Trial-level mean amplitudes and corresponding RPP values were concatenated across participants and analysed using a linear mixed-effects model, with subject included as a random intercept.

Additionally, given the reasonably large number of trials in PIT, we conducted time-frequency analyses to examine oscillatory dynamics during the processing of the Pavlovian and instrumental cues.

Data were then epoched between −500 and +2000 ms around the onset of the stimuli and baseline-corrected to −500–0 ms. Event-related spectral perturbations (ERSPs) were computed on the stimulus-locked epochs over a set of a *priori* regions of interest (ROIs), including the left pre-motor area (FC3; contralateral hemisphere to the hand participants used for performing the action) and frontal (Fz) area (Cavanagh et al., 2013; Finotti et al., 2025). Epochs in which premature responses were made were removed to ensure that the spectral estimates are not contaminated by motor-related activity.

ERSPs were computed with complex Morlet wavelets, applied across log-spaced frequencies from 3 to 30 Hz (80 bins). The wavelet parameters were set to two cycles at the lowest frequency with a scaling factor of 0.5, providing a balance between temporal and spectral resolution. Spectral estimates were derived across the full epoch window (−500 to 2000 ms relative to stimulus onset) and power values were baseline-normalised against the −500 to 0 ms interval. To improve stability, spectra were down-sampled to 200 time bins and zero-padded using a pad ratio of 2. For each subject, ERSPs were calculated separately for twelve bins corresponding to the full factorial combination of Pavlovian background valence (appetitive, aversive, neutral) and instrumental piggy bank conditions.

To test whether the cues influenced motor preparation, we extracted mean beta-band (13–30 Hz) power between 200 and 1500 ms post-stimulus from electrode FC3, as beta desynchronisation in this region is a well-established marker of motor readiness and action preparation (Pfurtscheller & da Silva, 1999; Kilavik et al., 2013). To assess cue-related cognitive control, we examined frontal theta activity (3–7 Hz, 50–500 ms post-stimulus at Fz). Frontal midline theta has been consistently linked to conflict monitoring, cognitive control, and the processing of motivationally salient events (Cavanagh & Frank, 2014; Cohen, 2014), making it a suitable index of Pavlovian influences on instrumental action.

### Results

#### Behaviour

A full-factorial RM-ANOVA (Background cues × Instrumental values) on key press rates showed a strong main effect of Pavlovian background, a main effect of instrumental value (reward per press), and a Background × Instrumental interaction (Table 2).

**Table 2:** Results of Full-Factorial Repeated-Measures ANOVA and Pairwise Comparisons for Key Press Rate.

| Effect | *df* | *F* or *t* | *p* | *M* (*SE*) |
| --- | --- | --- | --- | --- |
| *Main effects* |  |  |  |  |
| Pavlovian background | 6,372 | 44.23 | < .001 | — |
| Instrumental value (RPP) | 3,186 | 10.30 | < .001 | — |
| Background × Instrumental | 181,116 | 3.45 | < .001 | — |
| *Pairwise comparisons: Pavlovian background<sup>a</sup>* |  |  |  |  |
| *Background level means* |  |  |  |  |
| Aversive (−1) | — | — | — | 2.62 (0.31) |
| Aversive (−0.5) | — | — | — | 2.68 (0.30) |
| Aversive (−0.01) | — | — | — | 2.79 (0.30) |
| Neutral (0) | — | — | — | 4.72 (0.16) |
| Appetitive (0.01) | — | — | — | 5.23 (0.14) |
| Appetitive (0.5) | — | — | — | 5.17 (0.16) |
| Appetitive (1) | — | — | — | 5.30 (0.17) |
| *Selected contrasts* |  |  |  |  |
| Within aversive (all pairs) | — | *ts* < 2.03 | > .971 | — |
| Within appetitive (all pairs except below) | — | *ts* < 0.97 | = 1.000 | — |
| Appetitive 1-pound vs. 50-pence | 62 | 3.63 | .012 | — |
| All aversive vs. neutral & appetitive | — | *ts* > 6.04 | < .001 | — |
| *Pairwise comparisons: Instrumental value (RPP)<sup>a</sup>* |  |  |  |  |
| *RPP level means* |  |  |  |  |
| RPP 0.125 | — | — | — | 3.82 (0.15) |
| RPP 0.25 | — | — | — | 4.04 (0.14) |
| RPP 1 | — | — | — | 4.21 (0.15) |
| RPP 1.25 | — | — | — | 4.22 (0.16) |
| *Selected contrasts* |  |  |  |  |
| RPP 0.125 vs. 0.25, 1, and 1.25 | — | *ts* > 3.06 | < .020 | — |
| RPP 0.25 vs. 1.25 | 62 | 3.81 | .002 | — |
| All other contrasts | — | — | > .05 | — |
| *Interaction follow-up tests* |  |  |  |  |
| RPP effect, neutral & appetitive backgrounds | — | *F*s > 5.47 | < .001 | — |
| RPP effect, aversive backgrounds<sup>b</sup> | — | *F*s < 4.54 | > .004 | — |
| Background effect, all RPP levels | — | *F*s > 36.61 | < .001 | — |

Note. RPP = reward per press (pence per press). Means and standard errors are in presses per second.
<sup>a</sup> All pairwise *p*-values Bonferroni-corrected for multiple comparisons.
<sup>b</sup> No pairwise contrasts within aversive backgrounds survived Bonferroni correction.

Follow-up tests on the Pavlovian-instrumental interaction revealed that instrumental value effects were strong under neutral and appetitive backgrounds but they were weak or absent under aversive backgrounds. Conversely, background effects were robust at every instrumental level (Table 2).

To characterise individual differences in Pavlovian-instrumental transfer, we classified participants into Pavlovian and non-Pavlovian groups based on their key press behaviour. For each participant, we first calculated their mean key press rates (i.e., key presses per second) separately for trials with the most appetitive (associated with adding £1) and the most aversive (associated with losing £1) Pavlovian backgrounds. A PIT difference score was then computed by subtracting the mean rate in the most aversive from the most appetitive trials. Participants showing a Pavlovian influence (operationalised as a PIT difference score of at least one) were labelled as the Pavlovian group. Those with smaller or absent differences were labelled as the non-Pavlovian group. The PIT difference scores for the Pavlovian (*N* = 35) and non-Pavlovian (*N* = 28) groups are respectively 5.09 (*SE* = 0.28) and -0.31 (*SE* = 0.25). The group labels derived from this procedure are used in the subsequent ERSP analyses.

#### Readiness potential

The linear mixed effect model revealed that RPP did not significantly predict readiness potential amplitude (*F*(1, 7006) = 3.68, *p* = .055) with only a marginal trend toward more negative pre-motor activity at Cz for higher RPP values (readiness potential: *β* = −0.48 µV, SE = 0.25, 95% CI: [-0.97, 0.01]). While this suggests minimal modulation of readiness potentials by subjective reward values in the PIT task, the reduced range of RPP values (4 levels vs. 9 in the vigour task) may have limited sensitivity to detect this relationship despite the larger trial count. It is also possible that participants became less sensitive the instrumental values of stimuli in this paradigm when Pavlovian cues were also introduced.

#### ERSP

As there was no substantial difference in participants’ behaviour when comparing the background cues within valences, we combined trials with the same valence together to increase statistical power for the ERSP analysis. We performed a mixed-design ANOVA with Pavlovian background cues (appetitive, aversive or neutral) and Instrumental values (four RPP levels) as within-subjects factors and Group (Pavlovian or Non-Pavlovian) as a between-subjects factor.

##### Motor preparation (Beta suppression)

We found a significant main effect of Pavlovian cues, *F*(2, 732) = 6.12, *p* = .002. Follow-up comparisons showed that, appetitive backgrounds significantly suppressed the beta band activity relative to neutral background (*t*(62) = 4.03, *p < .*001) and aversive backgrounds (*t*(62) = 2.45, *p* = .017), see Figure 3C. There was no difference in beta band power between neutral and aversive backgrounds, *p* = .259.

There was also a significant main effect of Instrumental values, *F*(3, 732) = 3.09, *p* = .027. Specifically, the highest RPP condition suppressed the beta band activity significantly more than the lowest RPP condition (*t*(62) = 2.28, *p* =.026). No other pairwise contrasts were significant.

The main effect of group and the interaction effects were all non-significant, *Fs <* 1.81, *ps > .*095.

##### Motivational integration (Frontal theta)

We found no significant main effects of Instrumental values, *F*(3, 732) = 1.44, *p* = .230, or Group, *F <* 1, *p* = .574. No significant interactions were found, *Fs <* 1.71, *ps > .*164.

Although the main effect of Pavlovian cues did not reach significance, *F*(2, 732) = 2.96, *p* = .053, our planned pairwise comparisons showed that aversive cues elicited significantly stronger theta power than neutral cues (*t*(62) = 3.17, *p* = .002) and appetitive cues (*t*(62) = 2.28, *p* = .026), see Figure 3D. There was no significant difference between appetitive and neutral cues (*p* = .281). Further, exploratory analysis suggested this aversive enhancement (against neutral trials) was significant in the Pavlovian group (*p* = .018) but not in the Non-Pavlovian group (*p* = .065).

The frontal theta enhancement for aversive cues suggests recruitment of cognitive control networks that monitor potential conflicts between ongoing instrumental goals and aversive environmental signals.

#### Trial-by-trial neural-behavioural coupling

To test whether moment-to-moment neural fluctuations predict behavioural vigour, we conducted single-trial linear mixed-effects analyses relating oscillatory power to key-press rate. For each valid trial, beta power (13–30 Hz, 200–1500 ms) was extracted from FC3 as an index of motor preparation, and theta power (3–7 Hz, 50–500 ms) was extracted from Fz as an index of cognitive control. Power values were z-scored within participants. Models included Pavlovian condition and mean-centred instrumental value as covariates, with subject as a random intercept.

Beta power showed a significant negative relationship with key-press rate (*β* = −0.06, SE = 0.02, *t*(7522) = −3.17, *p* = .002), indicating that greater beta suppression predicted increased vigour. Frontal theta power was not significantly related to key-press rate (*β* = −0.01, SE = 0.02, *t*(7522) = −0.54, *p* = .588). This beta power-KPR coupling showed moderate test-retest reliability (*r* = 0.35), indicating that individual differences in the strength of motor preparatory beta suppression predicting behavioural vigour are partially stable across sessions.

## Experiment 4: Reversal learning task

The probabilistic reversal learning task probes the ability to detect and adapt to hidden changes in reward contingencies (Cools et al., 2002). Participants choose between two options whose reward contingencies periodically reverse. Optimal performance requires detecting reversals and rapidly adjusting behaviour. Reversal learning can involve inference over a latent, unobserved variable, and as such can involve more complex inference (Schlagenhauf et al., 2014; Mishchanchuk et al., 2024). Importantly, components of reversal learning appear to be selectively sensitive to dopamine and serotonin (Evers et al., 2005; den Ouden et al., 2013).

### Method

#### Stimuli

Cartoon squirrel images were sourced online and edited to serve as choice stimuli. Two squirrels were presented simultaneously, with one positioned on the left and one on the right side of the screen. Coin images depicting either high-value (£1) or low-value (1p) outcomes.

#### Procedure

Participants completed a probabilistic reversal learning task designed to examine learning in dynamically changing reward environments. The task was framed as a game in which participants chose between two squirrels, each with a bag of coins to share. Participants were instructed that one squirrel had higher-value coins (£1) while the other had lower-value coins (1p), and that the squirrels would “secretly switch bags” every few trials. The goal was to track which squirrel currently held the higher-value coins and maximise accumulated gains across the session. The task consisted of 40 (for pilot 1) and 30 (for pilot 2) blocks, with blocks alternating which squirrel held the optimal bag. Within each block, there were a maximum of 80 trials. A block could terminate early if participants accumulated a criterion number of optimal choices (criterion varied between 3 and 8 optimal choices across blocks).

As shown in Figure 1D, each trial started with a fixation cross presented for a random duration between 0.5 and 1 second, after which the two squirrel stimuli appeared. Participants had up to three seconds to make a choice by pressing the left or right arrow key. Upon selecting a squirrel, the outcome associated with the selection was revealed for one second, followed by the inter-trial interval of 500ms.

The reward contingencies varied across blocks. The first block was deterministic: choosing the optimal squirrel always resulted in the high-value outcome (£1), while choosing the suboptimal squirrel always yielded the low-value outcome (1p). Subsequent blocks introduced probabilistic feedback with gradually increasing uncertainty. In blocks 2-3, the optimal choice led to £1 on 90% of trials and 1p on 10% of trials. In blocks 4-5, this changed to 80% £1 and 20% 1p. From block 6 onwards, the contingencies stabilised at 70% £1 and 30% 1p for optimal choices. This probabilistic structure created occasional surprising outcomes even when participants selected the correct option, allowing us to distinguish between surprise arising from probabilistic noise within stable contingencies versus surprise arising from actual contingency changes at block boundaries.

#### EEG pre-processing and analysis

We epoched the data time-locked to the onset of outcome images, using the interval be-tween −100 and 0 ms before the onset of outcomes as the baseline for correction.

We used a mass-univariate GLM to analyse trial-wise EEG. For each participant and time-locked event, an OLS GLM with an intercept was fit at every electrode and time sample using trial-level behavioural regressors z-scored within participant. To examine how accumulated learning experience modulates outcome processing, we constructed a GLM including: (1) accumulated optimal feedback (z-scored count of optimal outcomes received since the last reversal), (2) outcome valence (optimal coded as 1, suboptimal coded as -1), (3) reaction time (z-scored), and (4) the interaction between accumulated optimal feedback and outcome valence. This analysis was restricted to stay trials (where participants repeated their previous choice) to isolate learning-related signals from choice-switching processes. The interaction term allowed us to test whether accumulated experience differently modulated processing of optimal versus suboptimal outcomes.

In addition, to directly test whether the brain differentiates between surprise types, we compared the neural activity in switch trials (a true contingency reversal, irrespective of the participant’s choice on that trial; unstable context) with trials with probabilistic uncommon feedback (in stable context). Surprise type (stable surprise coded as 1, unstable surprise coded as −1) and reaction time were included in this GLM.

### Results

#### Behaviour

To examine response strategies following feedback, we computed win-stay probability (the proportion of trials on which participants repeated their previous choice following optimal feedback) and lose-shift probability (the proportion of trials on which participants switched following suboptimal feedback).

Participants demonstrated strong win-stay behaviour across both stable (0.86) and unstable (i.e., switch trials; 0.87) contexts, indicating robust exploitation of rewarding choices. Lose-shift probabilities were 0.70 and 0.74 in stable and unstable contexts, respectively. Figure 4A shows the learning curve around reversals.

**Figure 4:**
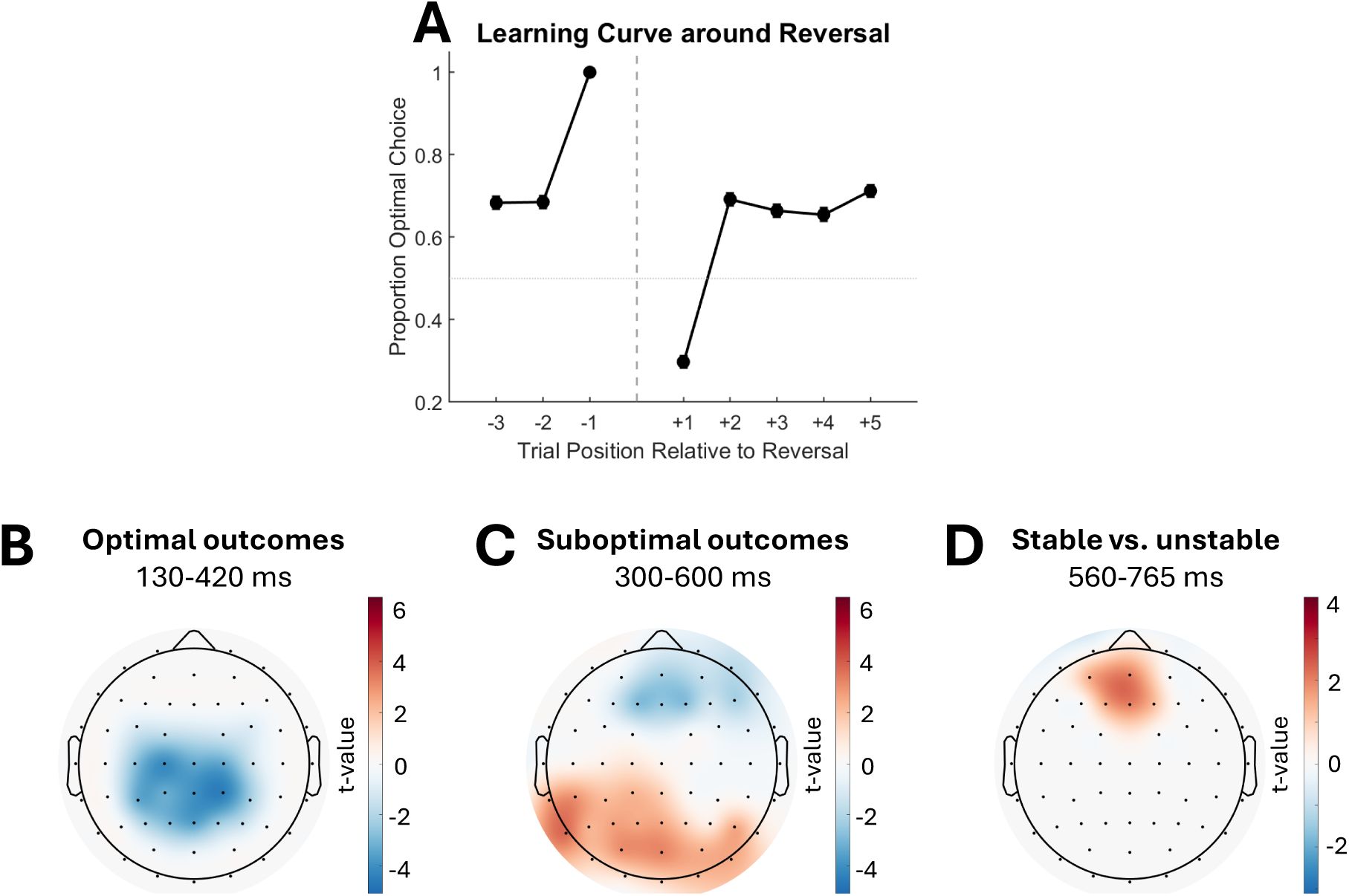
(A) Learning curves around reversals. (B) Scalp topographies showing significance-thresholded effect of accumulated optimal feedback on neural responses to optimal outcomes, (C) significance-thresholded effect of accumulated optimal feedback on neural responses to suboptimal outcomes, and (D) the contrast between surprising outcomes in stable vs. unstable contexts.

#### Neural signatures of accumulated learning during reversal

To investigate how accumulated learning experience shapes outcome processing, we examined EEG responses as a function of accumulated optimal feedback. For optimal outcomes, increased accumulated optimal feedback was associated with enhanced centro-parietal negativity spanning 130-420 ms post-outcome. This effect was broadly distributed across central and parietal electrodes (see Figure 4B), indicating that as participants accumulated more evidence about the optimal stimulus, confirmatory feedback elicited increasingly negative potentials in this region. This sustained negativity may reflect efficient predictive processing, where expected outcomes require less neural resources for evaluation as certainty increases. This beta coefficient (averaged over CP2) of the effect of accumulated learning negatively correlated with the beta coefficient (averaged over CP2) of the effect of trial progression from the PILT (r = -0.34), suggesting that a common centro-parietal learning signal underlies both tasks.

Conversely, for suboptimal outcomes, accumulated learning produced a markedly different pattern. Higher accumulated optimal feedback predicted a frontal negativity (see Figure 4C) and a simultaneous parietal-occipital positivity (P300) between 300 and 600 ms. This dipolar pattern suggests that as participants develop stronger expectations about optimal choices, violations of these expectations trigger a robust evaluative response combining frontal error detection with posterior evaluative processes to unexpected outcomes.

#### Context-dependent modulation of surprise processing

We next examined whether the brain differentiates between surprise occurring in stable versus unstable contexts. Comparing neural responses to unexpected outcomes revealed that surprise in stable contexts (i.e., uncommon probabilistic suboptimal feedback) elicited a frontal positivity (see Figure 4D) relative to surprise in unstable contexts (i.e., switch trials), beginning approximately 560 ms post-outcome.

This context-dependent modulation indicates that the brain adaptively calibrates neural responses to surprises based on environmental volatility. Specifically, surprising outcomes at switch trials elicit attenuated frontal responses compared to surprising outcomes during stable contingencies, when expectations may be well-established. This is intriguing because, at the moment of outcome delivery, participants could not have known whether it was a switch trial or not. Perhaps, the reduced neural response to switch trials reflects an implicit volatility tracking: in this task, switch trials usually occurred after a certain level of adaptation to the previous contingency, and the brain may have learned that such extended periods of stability tend to precede an upcoming reversal. After extended learning or adaptation, the brain may engage a conservative updating mechanism that anticipates change and dampens responses to surprising outcomes following prolonged periods of consistent feedback. In contrast, probabilistic errors that occur intermittently throughout stable periods, without being preceded by this same extended run of consistent feedback, are not treated as signalling an upcoming reversal, and likely maintain their capacity to drive frontal error signals, which may reflect ongoing learning.

## Experiment 5: Reinforcement learning under working memory load

The working memory load RL task aims to dissociate incremental RL from rapid inference supported by working memory (Collins & Frank, 2012, 2018). This task is akin to the PILT task, but with two key changes. First, the task involves presentation of one stimulus and choice between three actions, rather than the identification of a stimulus. This shifts memory load from recognition towards encoding of abstract, non-observed associations. Second, the task manipulates the delay between presentations of the same stimulus, enabling a parametric assessment of the effect of delay on the strength with which reward outcomes are associated with stimuli. At short stimulus delays, correct responses can be retrieved directly from active WM maintenance. As delays increase, WM traces decay and participants must rely on gradual RL (Westbrook et al., 2025; Collins, 2026). These mechanisms are thought to recruit separable neural systems, making the task useful for assessing their relative contributions and identifying selective pharmacological effects on each system.

### Method

#### Stimuli

Fourteen new object images were chosen from the picture database used in Experiment 1 and were perceptually matched.

#### Procedure

The working memory task examined how reinforcement learning performance varies under different working memory loads. In each trial, participants were presented with a single card stimulus at the centre of the screen. Participants had up to 3 s to choose one of three response keys (left, up, or right arrow) to “flip” the card and reveal the associated outcome (1p, 50p, or £1). For each card, one key was consistently associated with a higher coin outcome, while the other two yielded lower outcomes (i.e., deterministic feedback). Upon a selection, the outcome and the selected key icon are presented for one second (Figure 1E).

Working memory was manipulated by varying the delay between repetitions of each stimulus. Specifically, a short delay (delay 1) created low working memory load, as the stimulus reappeared immediately. Intermediate delays (delay 2-4) imposed moderate working memory load, requiring limited maintenance of the stimulus-action mapping across a few intervening trials. Long delays (delay *>* 4) imposed high working memory load, as participants had to retrieve the correct response after several other stimuli.

There were in total 108 trials, and there was no break in the task.

#### EEG pre-processing and analysis

We epoched EEG time-locked to the onset of outcome images (time range: from -200 to 800 ms) or the onset of object images (time range: from -200 to 600 ms), using the −200-0 ms interval as baseline. The EEG analyses followed the same pipeline as Experiment 1. Specifically, for each participant, an OLS GLM with an intercept was fit at every electrode and time sample, modelling the outcome-locked amplitudes as a function of reward magnitude, and the stimulus-locked amplitudes as a function of stimulus repetition. Additionally, we ran single-predictor (reward magnitude or stimulus repetition) GLMs separately for each delay condition: short (delay 1), intermediate (delay 2-4), and long (delay *>* 4). *β*-coefficients at all electrodes and time points yielded 3-D *β* -maps. Group-level inference tested *β* against zero using two-sided sign-flip permutations with TFCE across channels and time points (5,000 permutations; family-wise error controlled at *α* = .05 via the max-TFCE statistic).

### Results

#### Behaviour

A multiple logistic regression with delay, stimulus repetition, and their interaction revealed that participants’ performance was influenced by both stimulus repetition and delay. Participants were more likely to select the correct choice with increased stimulus appearances (*β* = 0.34, z = 13.09, *p < .*001; OR = 1.40, 95% CI: [1.33, 1.47]) and longer delays since last appearance (*β* = 0.11, z = 3.70, *p < .*001; OR = 1.11, 95% CI: [1.05, 1.17]). A significant interaction between appearance and delay (*β* = -0.02, z = -4.49, *p < .*001; OR = 0.98, 95% CI: [0.97, 0.99]) indicated that the benefit of repetition diminished as delay increased. Learning curves were plotted separately for the three delay conditions in Figure 5A.

**Figure 5:**
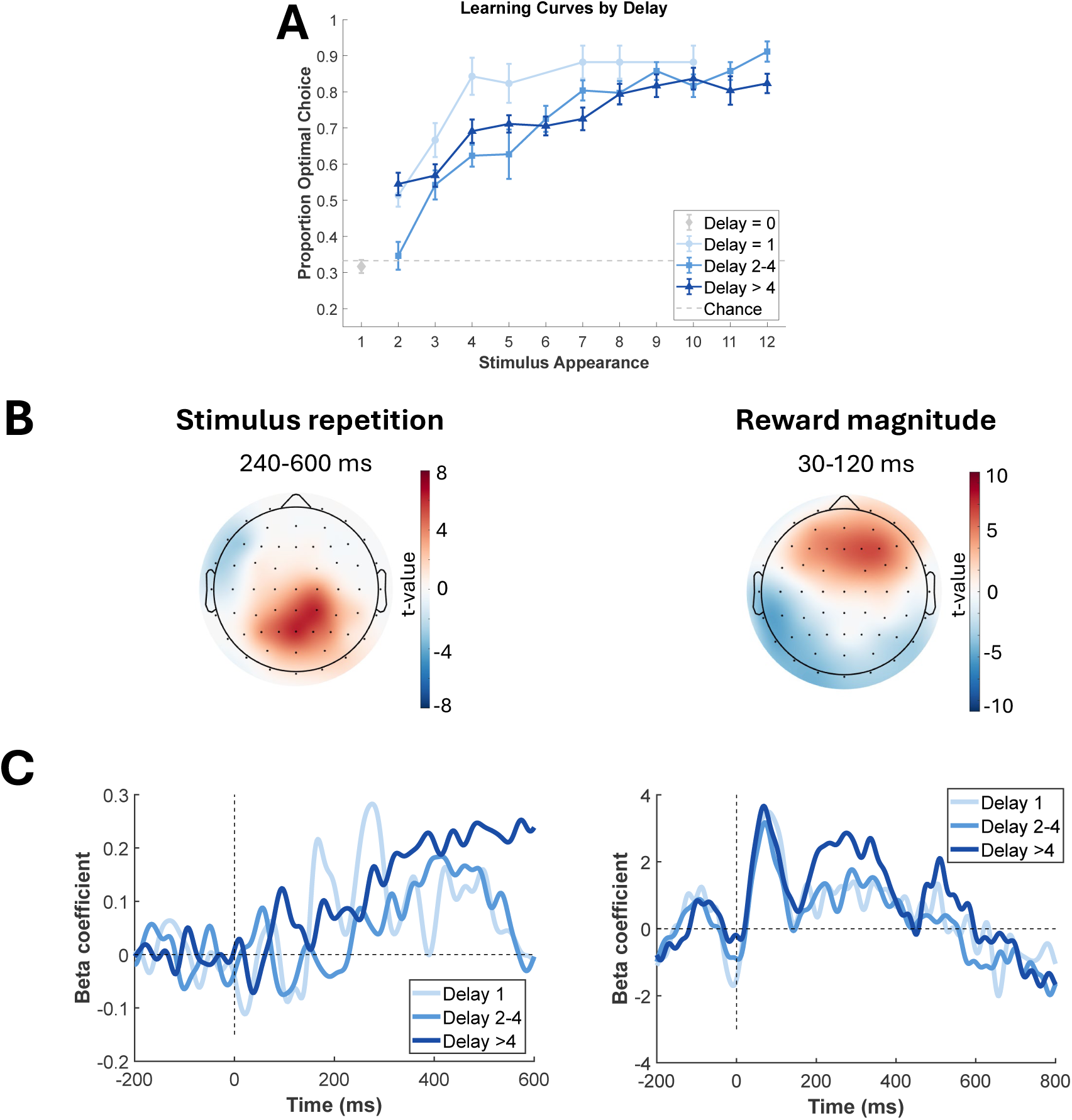
(A) Learning curves by delay levels. (B) Scalp topographies plotting significance-thresholded average regression weights for stimulus repetition effects on stimulus-locked EEG and reward magnitude effects on outcome-locked EEG. (C) Time course of the average beta coefficients over electrodes showing maximal effects of stimulus repetition (left) and reward magnitude (right), separately for three delay conditions.

#### EEG

In examining the effect of stimulus repetition on stimulus-locked EEG signals, a significant centro-parietal cluster (spatial peak: Pz) from 240-600 ms post-stimulus (temporal peak: 414 ms) was found, whereby later trials (more repetitions of the same stimulus) were associated with a stronger P300 than earlier trials (less repetitions of the same stimulus), *t*(46) = 7.62, TFCE-corrected *p < .*05 (Figure 5B). This effect may reflect stimulus learning, which is similar to the trial-progression marker (i.e., enhanced P300 for later trials) found in the PILT. Indeed, the beta coefficient (averaged over CP2) for the effect of trial progression from the PILT positively correlated with the beta coefficient (averaged over Pz) for the effect of stimulus repetition in this task (r = 0.30).

As our a *priori* comparisons of interest, single-predictor GLMs were conducted for each delay condition. We found that this centroparietal positivity towards later trials had an earlier onset when the repetition immediately followed the last stimulus appearance (onset at around 240ms post-stimulus). However, the onset of the repetition effect was much later when the delay was larger (i.e., delay *>* 4; onset after 300ms post-stimulus), see Figure 5C. Interestingly, the effect of stimulus repetition was not significant for the delay 2-4 condition.

Two time windows were found to show significant effects of the reward magnitude on EEG time-locked to outcomes. Specifically, between 30 and 120ms after the onset of outcomes, there was a frontal positivity for trials with larger rewards (spatial peak: F4, temporal peak: 75ms; *t*(43) = 10.59, TFCE-corrected *p < .*05; Figure 5B). In the same time window, the signals over posterior electrodes were more negative for larger than smaller rewards. Similarly, at a later time window (200-500ms), a frontal positivity (*ts >* 2.48, TFCE-corrected *ps < .*05) and a parietal negativity (*ts >* 2.02, TFCE-corrected *ps < .*05) were found for larger rewards, compared to smaller rewards. As shown in Figure 5C, while the early effect of reward magnitude was found in all three delays conditions, the later effect of PE was only observed when the working memory load was higher.

## Discussion

In the present study, we developed a multi-task EEG battery spanning five reinforcement learning paradigms, each targeting a distinct RL sub-component. The probabilistic instrumental learning task, designed to measure incremental value learning from probabilistic feedback, elicited a centroparietal P300 scaling with trial progression. The working memory-RL task, designed to dissociate rapid WM-supported learning from slower incremental RL, showed a closely analogous P300 repetition effect with a delay-dependent onset. The vigour task, targeting motivational modulation of action under varying reward rates, revealed a readiness potential at Cz that scaled with reward-per-press, indicating value-dependent motor preparation. The PIT task, designed to examine how Pavlovian cues modulate ongoing instrumental behaviour, revealed dissociable markers of motor preparation and conflict processing. Specifically, pre-motor beta suppression over FC3 was sensitive to both Pavlovian valence and instrumental value, while frontal theta was selectively enhanced by aversive cues. Finally, the reversal learning task, probing behavioural adaptation to hidden changes in reward contingencies, produced dissociable centroparietal signatures for confirmatory and disconfirmatory outcomes, and a late frontal positivity distinguishing surprise in stable versus volatile contexts. Taken together, these findings identify EEG markers sensitive to distinct RL sub-components, whose largely independent patterns across tasks suggest that the battery tracks separable reinforcement learning processes. This is critical as previous studies have at times found that different RL and decision-making tasks may be dominated by a single underlying ‘task acuity’ dimension (Moutoussis et al., 2021), and such a single underlying dimension would likely undermine attempts at identifying selective behavioural correlates of underlying neuromodulatory systems (Mkrtchian et al., 2025).

### Neural markers of incremental value learning

A centro-parietal P300 that scaled with trial progression emerged as the most consistent neural signature of learning across the battery. In the blocked PILT (Experiment 1a), stimulus-locked amplitudes at CP2 increased over the course of each block, and this pattern was replicated in the mixed-valence version (Experiment 1b). A strikingly similar effect appeared in the working memory reinforcement learning task (Experiment 5), where the centroparietal P300 scaled with the number of times a stimulus had been encountered. The convergence of this effect across three independent paradigms suggests that the P300 indexes a domain-general learning signal that tracks the accumulation of stimulus-outcome knowledge regardless of task structure and whether the stimulus set is large or small.

This interpretation aligns with the context-updating account of the P300 (Donchin & Coles, 1988), according to which the component reflects the revision of an internal model in response to task-relevant information. As learning progresses, each stimulus presentation carries more accumulated associative information, and the P300 may reflect the integration of this accruing evidence into an updated value representation (Twomey et al., 2015). The positive cross-task correlation between the PILT trial-progression betas and the WM-RL stimulus-repetition betas provides support for this claim: participants who showed a stronger P300 learning signal in one task tended to do so in the other. This cross-task correlation is notable given that PILT and WM-RL differ substantially in stimulus set size, feedback determinism, and working memory demands, suggesting the shared variance reflects a genuine individual-difference trait in incremental learning rather than task-specific factors.

Interestingly, the learning-related P300 observed in the reversal learning task (Experiment 4) showed a negative relationship with the PILT marker. In the reversal task, accumulated optimal feedback was associated with an increasingly negative centroparietal deflection for confirmatory outcomes (i.e., suppressed P300). This apparent contradiction is, however, consistent with the differing computational demands of the two paradigms. In the PILT and WM-RL tasks, later trials carry progressively more value-relevant information, and the P300 tracks this growing informational content. In the reversal task, by contrast, accumulated confirmatory evidence reflects increasing certainty about the current contingency. As expected, outcomes become more predictable and less evaluative processing is required, yielding reduced (more negative) centroparietal amplitudes to confirmatory feedback.

### Neural markers of outcome evaluation

Outcome-locked analyses revealed a sustained frontocentral negativity for rewards relative to losses in both PILT versions. This pattern resembles a loss-related positivity, likely corresponding to the error-related positivity (Pe; Falkenstein et al., 2000; Ridderinkhof et al., 2009). The Pe is a sustained centroparietal positivity associated with the conscious appraisal of unfavourable outcomes, and its enhancement for losses relative to rewards in the current data is consistent with heightened evaluative processing when feedback signals a suboptimal choice (Ridderinkhof et al., 2009).

A more nuanced picture emerged when we examined how outcome magnitude modulated ERP components within each valence context, as this pattern was strikingly dependent on task structure. In the blocked design (Experiment 1a), magnitude effects were asymmetric across valence: smaller rewards enhanced the P300, whereas larger losses enhanced the P100 over posterior electrodes. These two effects likely reflect distinct mechanisms. The enhanced P300 for smaller rewards is consistent with a surprise or oddball effect within reward blocks, where high-value outcomes (50p and £1) are frequent, the occasional 1p outcome is unexpected, and the P300 may track this relative infrequency rather than outcome value *per se*. The enhanced P100 for larger losses, by contrast, suggests rapid attentional capture by the most aversive outcome in the set, consistent with evidence that negatively-valenced stimuli receive prioritised early visual processing (Pourtois et al., 2013).

In the mixed-valence design (Experiment 1b), this asymmetry disappeared. The P100 was enhanced for larger outcomes in both valence directions, while the P300 enhancement for smaller rewards was absent altogether. When valence is blocked in Experiment 1a, participants can anticipate the evaluative framework for each trial, for example, seeing a 1-penny coin immediately indicates that this is the worst outcome (in fact, the punishment) in this block and that the better outcome for this block will be a 1-pound coin. This strategic, context-specific processing is indeed reflected in the behavioural data with an early plateau and overall higher accuracy in Experiment 1a, compared to 1b.

When valence is unpredictable on a trial-by-trial basis, however, participants cannot adopt such context-specific strategies. Instead, the early visual system appears to encode outcome magnitude symmetrically across both valence directions, as reflected in the bilateral P100 sensitivity. The disappearance of the P300 oddball effect for smaller rewards is likewise expected because the 1p outcomes are no longer infrequent in this design, removing the surprise that drove the P300 in the blocked version.

A feedback-related negativity (FRN) to larger relative to smaller losses was observed over frontal electrodes in the blocked PILT. Similarly, a FRN was also found for the suboptimal outcomes in the WM-RL task. The FRN has been widely linked to rapid negative outcome evaluation, possibly generated by the anterior cingulate cortex in response to worse-than-expected outcomes (Gehring & Willoughby, 2002; Holroyd & Coles, 2002). Its restriction to punishment blocks in Experiment 1a is consistent with the view that the FRN is most robustly elicited during punishment learning. This explanation isn’t incompatible with its presence in the WM-RL task, however. It is likely that the FRN may be most suitable as a biomarker for negative outcome evaluation in paradigms where the learning context is unambiguous (i.e., only comprising loss outcomes or fully deterministic).

### Dissociating working memory and incremental reinforcement learning

The WM-RL task was added to the battery for us to potentially dissociate rapid, WM-supported learning from slower, incremental RL. This manipulation seems to be effective both behaviourally, whereby the benefit of stimulus repetition diminished as delay increased, and neurally, where it modulated both the timing and the presence of key ERP effects.

Specifically, for stimulus-locked activity, the P300 repetition effect showed an earlier onset at short delays compared to long delays. This latency shift is consistent with the idea that short-delay learning likely relies on active WM maintenance, which enables rapid retrieval and early neural evidence accumulation, whereas long-delay learning requires slower, RL-dependent retrieval that shifts the onset of the P300 to a later time point (e.g., Collins & Frank, 2012).

The outcome-locked analyses revealed what may be the most informative dissociation in this task. Reward magnitude modulated EEG amplitudes in two distinct time windows: an early frontal positivity with concurrent posterior negativity (30-120ms), and a later frontal positivity with parietal negativity (after 200ms). Critically, while the early magnitude effect was present across all delay conditions, the late effect was observed only under higher working memory load. This dissociation maps directly onto the theoretical framework proposed by Collins and Frank (2012), in which WM and RL operate as separate systems. When WM resources are available (short delays), participants can rapidly infer the correct response from a single or few experiences, a process that may not engage the incremental learning computations reflected in later ERP components. The early magnitude effect, present at all delays, likely reflects a more automatic perceptual processing of outcome values. However, when WM load is high (long delays), participants must rely on trial-by-trial reinforcement learning, and a later effect of higher-level outcome evaluation emerges.

These learning-related markers have direct relevance for pharmacological research targeting depression. The loss-related Pe and the FRN, both preferentially elicited by negative outcomes, may be particularly sensitive to serotonergic interventions, given converging evidence that serotonin modulates aversive outcome processing and punishment learning (Cools et al., 2008; Colwell et al., 2024; Dayan & Huys, 2009; Michely et al., 2022). The WM-RL dissociation offers further specificity: because WM-dependent and RL-dependent learning recruit separable prefrontal and striatal-dopaminergic systems respectively (Collins & Frank, 2012; Collins et al., 2017), neuromodulators that primarily affect prefrontal function may selectively modulate learning at short delays, whereas dopaminergic agents acting on striatal circuits may preferentially alter the late outcome evaluation signal that emerges under high working memory load.

### Neural signatures of cognitive flexibility

The reversal learning task extended the learning framework by examining how the brain processes outcomes in a dynamically changing environment. Unlike the PILT and WM-RL tasks, where contingencies are stable within blocks, the reversal task required participants to detect hidden changes and rapidly adjust behaviour. As a form of cognitive flexibility, it is thought to depend on both dopamine-driven inference about hidden states (Costa et al., 2015; Izquierdo et al., 2017) and on serotonin (Evers et al., 2005; Clarke et al., 2004, 2005)..

Accumulated optimal feedback produced two dissociable outcome-processing signatures depending on whether the feedback confirmed or violated current expectations. Specifically, as participants develop stronger expectations about the current contingency, unexpected outcomes trigger a frontal error detection and a posterior context updating. While the frontal component is consistent with the established role of medial frontal cortex in signalling outcomes that deviate from expectations (i.e., the error-related negativity; Cavanagh & Frank, 2014; Ullsperger et al., 2014), the posterior P300 likely reflects the need to revise the internal model of the current reward structure.

We also explored whether the brain differentiates between surprise occurring in stable versus unstable contexts, by comparing neural responses to unexpected suboptimal outcomes during stable contingencies (probabilistic errors) with those at reversal points (switch trials). This comparison revealed a late frontal positivity for stable-context surprises relative to unstable-context surprises. This finding is broadly consistent with the idea that the brain may calibrate its response to surprising outcomes based on the recent history of environmental stability (Behrens et al., 2007; Nassar et al., 2019). While computational models (Browning et al., 2015; Mathys et al., 2011) demonstrate that learning rates should increase during volatility, our findings suggest that a neural signature of expectancy violation (i.e., the frontal positivity) may actually be maximal when a stable, and potentially highly-weighted prediction is breached. Specifically, after prolonged consistent feedback, sporadic probabilistic errors may maintain a high capacity to drive frontal error signals due to the high contrast with established expectations. Conversely, in volatile contexts, the brain may engage a more conservative evaluative mechanism for individual outcomes, reflecting an implicit tracking of environmental instability where single surprises are expected and thus elicit attenuated neural responses.

### Neural markers of motivational vigour

The vigour and PIT tasks were designed to capture how the value of stimuli and the emotional significance of environmental cues jointly shape the motivation to act. We focused on three distinct neural markers: the readiness potential (RP) and pre-motor beta suppression that track motor preparation, and frontal theta that indexes complementary aspects of motivational and conflict processing.

In the vigour task, the RP at Cz overall scaled with reward-per-press, such that higher-value actions were preceded by more negative pre-motor activity. This finding is consistent with the view that the RP reflects the motivational intensity with which an action is initiated (Deecke, 1996; Fried et al., 2011). Within the framework of average reward rate theories of vigour (Niv et al., 2007), stronger motor preparation for high-value actions may reflect a dopaminergic computation of net action utility, whereby the tonic dopamine signal (which encodes opportunity cost) biases the motor system toward more energetic responding when the expected return is high. This value-dependent modulation of the RP provides a neural counterpart to the behavioural finding that key-press rates increased with reward-per-press, and suggests that motivational influences on action are already present at the level of motor preparation, prior to movement onset.

Time-frequency analysis in the PIT task revealed that pre-motor beta desynchronisation over FC3 was modulated by both Pavlovian cues and instrumental values. Beta desynchronisation in lateral premotor regions is a well-established index of motor readiness and action preparation (Pfurtscheller & da Silva, 1999; Kilavik et al., 2013). Appetitive backgrounds produced significantly greater beta suppression than neutral and aversive backgrounds, consistent with a Pavlovian approach bias that primes the motor system for action in rewarding contexts (Huys et al., 2011). Similarly, the highest RPP stimulus produced greater suppression than the lowest RPP stimulus, showing that the beta power change also reflected the instrumental values of actions.

Importantly, trial-by-trial beta power negatively predicted key-press rate, showing that greater beta suppression on a given trial may have translated into more vigorous responding. The combination of condition-level sensitivity and trial-level behavioural prediction positions pre-motor beta suppression as a promising biomarker for motivational vigour.

Lastly, the frontal theta indexes Pavlovian conflict well in our paradigm. Specifically, aversive Pavlovian backgrounds elicited stronger theta power than both neutral and appetitive cues. Frontal theta has been consistently linked to conflict monitoring and cognitive control (Cavanagh & Frank, 2014; Cohen, 2014), and its selective enhancement under aversive cues likely reflects a Pavlovian-instrumental conflict. That is, the Pavlovian impulse to inhibit action in the face of aversive cues (i.e., previously associated with losing coins) and the ongoing instrumental goal of pressing to earn rewards formed some level of conflict. The frontal theta band activity was responsive to this conflict. Indeed, the anterior cingulate cortex (ACC), a key node in conflict monitoring and adaptive control (Botvinick et al., 2001; Shenhav et al., 2013), is widely regarded as the primary cortical generator of scalp-recorded midline frontal theta (Cavanagh & Shackman, 2015; Ishii et al., 1999). The aversive theta enhancement observed here is therefore consistent with ACC-mediated detection of Pavlovian-instrumental conflict, whereby the brain processes the misalignment between cue-evoked inhibitory tendencies and task-relevant approach behaviour.

In summary, these results suggest a functional dissociation within the neural mechanisms underlying PIT. While frontal theta power indexes the cognitive effort required to resolve competing motivational signals, beta desynchronisation over the pre-motor cortex serves as a proxy for the final motivational “push” towards action.

From a clinical perspective, these markers may also help address distinct features of depression. Motivational deficits such as reduced vigour, psychomotor slowing, and anhedonia are among the core symptoms of depression and are thought to reflect disrupted dopaminergic encoding of action value and opportunity cost (Husain & Roiser, 2018; Treadway & Zald, 2011). The RP and beta suppression markers may be sensitive to dopaminergic interventions that restore the value-based modulation of motor preparation. Meanwhile, pharmacological interventions targeting serotonergic systems, which are implicated in both Pavlovian inhibition and the processing of aversive outcomes (Cools et al., 2008; Dayan & Huys, 2009), may selectively modulate this theta-indexed conflict signal while leaving the dopaminergic motor preparation pathway relatively intact.

### Cross-task convergent and divergent validity

A central aim of this battery is to provide multiple neural markers that can be selectively targeted by pharmacological interventions acting on distinct neurotransmitter systems. This goal requires that the battery captures both shared and independent processes. Therefore, we hoped to identify markers that converge where the underlying computation is common, and those diverge where the tasks tap into separable neural mechanisms.

On the convergent side, the P300 learning signal showed consistent cross-task relationships. As discussed earlier, the centro-parietal P300 likely captures a common learning-related process. This convergence is encouraging because it implies that a pharmacological effect on this mechanism should be detectable across multiple tasks in the battery, increasing statistical power and interpretive confidence. A complementary form of convergent evidence came from the representational similarity analysis in the PILT. The shift in neural representational geometry following learning, whereby post-learning EEG patterns in passive viewing aligned more closely with learned stimulus values, demonstrates that the PILT drives a lasting change in stimulus representation. This finding, to our knowledge the first application of RSA to passive viewing pre/post instrumental learning in an EEG battery context, suggests that value learning leaves a representational signature that could be tracked pharmacologically independently of task performance.

On the divergent side, the vigour and PIT tasks targeted motivational processes thought to depend on tonic dopaminergic signalling (Niv et al., 2007), whereas the PILT outcome evaluation markers (Pe, FRN) index feedback processing linked to serotonergic function (Cools et al., 2008; Michely et al., 2022). The beta-theta dissociation within the PIT task itself already demonstrates divergent validity at the within-task level: beta suppression predicted trial-by-trial vigour while frontal theta did not, indicating that motor preparation and conflict monitoring are separable processes even when measured in the same paradigm.

The test-retest reliability data, though currently available only for a subset of markers, provide a preliminary basis for evaluating the suitability of these measures as biomarkers. For the mixed-valence PILT, most neural markers showed moderate reliability across sessions separated by 11-19 days (Pearson’s r ranging from 0.46 to 0.61). In the PIT task, the trial-level coupling between beta power and key-press rate showed moderate reliability (r = 0.35). These values fall within the range commonly reported for task-evoked ERP and oscillatory measures, which typically exhibit lower reliability than raw amplitude or power estimates (e.g., Cassidy et al., 2012). Given that trial-level coupling coefficients incorporate variance from both neural and behavioural measures, some attenuation of stability is expected. Nevertheless, moderate reliability indicates that a meaningful proportion of individual differences in these neural signatures is preserved over time.

Our findings provide an initial framework for understanding convergence and divergence across the included tasks, but they do not constitute a comprehensive validation of the measures. Future work is needed to include external cognitive or clinical measures that would allow us to test criterion validity. For example, whether our motivational markers predict self-reported anhedonia should be tested. Second, while the current model-free analytical approach offers transparency and avoids computational assumptions, it may have underestimated the true similarity or dissimilarity between tasks, as the empirical proxies used (e.g., accumulated feedback) likely captured somewhat different slices of the underlying computation in each task. Future work incorporating computational modelling as a complementary approach could provide a more formal decomposition of shared and unique variance across tasks.

Finally, it is important to emphasise that the battery is not intended as a set of interchangeable tasks, but as an integrated framework in which each paradigm contributes a distinct component of reinforcement learning. Across the battery, the tasks span incremental learning from feedback, motivational vigour under varying action costs, Pavlovian modulation of instrumental behaviour, behavioural flexibility under contingency change, and the interaction between working memory and incremental learning. These components are thought to rely on partially dissociable neuromodulatory mechanisms, and therefore cannot be captured by any single task or neural marker in isolation.

From an applied perspective, this structure allows us to test whether a pharmacological manipulation produces broad effects across reinforcement learning systems or selectively modulates a specific sub-component. This distinction is clinically relevant, given evidence that serotonergic and dopaminergic agents exert dissociable influences on different aspects of reinforcement learning (Mkrtchian et al., 2025).

## Conclusion

The present study establishes a multi-task EEG battery that yields dissociable neural biomarkers of incremental learning, motivational vigour, Pavlovian-instrumental transfer, cognitive flexibility, and the interplay between working memory and incremental learning. Across the learning paradigms, a centroparietal P300 seems to be a consistent marker of learning, and outcome evaluation components showed systematic sensitivity to valence, magnitude, and task structure. In our motivational vigour tasks, readiness potential and beta-band suppression tracked the values of actions, and frontal theta indexed Pavlovian conflict. The convergent validity observed for the P300 learning signal, together with the largely independent patterns across other markers, supports the battery’s capacity to detect selective pharmacological effects on distinct neural systems. Future work will combine these neural data with specific computational models to better identify the neural representation of specific computational processes. Overall, these findings provide a foundation for deploying this battery in studies aimed at understanding how different interventions shape the neural computations underlying learning, motivation, and adaptive behaviour.

## Declarations

### Ethics approval

This study was performed in line with the principles of the Declaration of Helsinki. The study was approved by the UCL Research Ethics Committee (project ID: 1273).

### Consent to participate

Informed consent was obtained from all individual participants included in the study.

### Consent for publication

Not applicable.

## Acknowledgments

We acknowledge support by the members of the Applied Computational Psychiatry Lab.

## Open Practices Statement

Data or materials for the experiments are available upon request, and none of the experiments was preregistered.

